# lra: the Long Read Aligner for Sequences and Contigs

**DOI:** 10.1101/2020.11.15.383273

**Authors:** Jingwen Ren, Mark JP Chaisson

## Abstract

**Motivation:** It is computationally challenging to detect variation by aligning long reads from single-molecule sequencing (SMS) instruments, or megabase-scale contigs from SMS assemblies. One approach to efficiently align long sequences is sparse dynamic programming (SDP), where exact matches are found between the sequence and the genome, and optimal chains of matches are found representing a rough alignment. Sequence variation is more accurately modeled when alignments are scored with a gap penalty that is a convex function of the gap length. Because previous implementations of SDP used a linear-cost gap function that does not accurately model variation, and implementations of alignment that have a convex gap penalty are either inefficient or use heuristics, we developed a method, lra, that uses SDP with a convex-cost gap penalty. We use lra to align long-read sequences from PacBio and Oxford Nanopore (ONT) instruments as well as *de novo* assembly contigs.

**Results:** Across all data types, the runtime of lra is between 52-168% of the state of the art aligner minimap2 when generating SAM alignment, and 9-15% of an alternative method, ngmlr. This alignment approach may be used to provide additional evidence of SV calls in PacBio datasets, and an increase in sensitivity and specificity on ONT data with current SV detection algorithms. The number of calls discovered using pbsv with lra alignments are within 98.3-98.6% of calls made from minimap2 alignments on the same data, and give a nominal 0.2-0.4% increase in F1 score by Truvari analysis. On ONT data with SV called using Sniffles, the number of calls made from lra alignments is 3% greater than minimap2-based calls, and 30% greater than ngmlr based calls, with a 4.6-5.5% increase in Truvari F1 score. When applied to calling variation from de novo assembly contigs, there is a 5.8% increase in SV calls compared to minimap2+paftools, with a 4.3% increase in Truvari F1 score.

**Availability and implementation:** Available in bioconda: https://anaconda.org/bioconda/lra and github: https://github.com/ChaissonLab/LRA

## 1 Introduction

Studies of genetic variation often begin by aligning sequences from a sample back to a reference genome, and inferring variation as differences in the alignment. Long read, single molecule sequencing (LRS) is becoming established as a routine approach for sequencing genomes. The two technologies that produce LRS technologies, Pacific Biosciences (PacBio) and Oxford Nanopore (ONT) generate reads over 50kb at error rate 15% or less. Aligning these sequences is a computationally challenging task for which several methods are available including minimap2, ngmlr, and BLASR (Li, 2018; Sedlazeck *et al.*, 2018; Chaisson and Tesler, 2012). They are demonstrated to be quite fast and accurate, but have limitations, particularly when there are large sequence differences between the read and the reference. This problem is amplified in complex, repetitive regions such as variable-number tandem repeats, that only make up 6% of the human genome, but account for nearly 70% of observed structural variation: insertions and deletions at least 50 bases (SV), and in larger SV (Rowell *et al.*, 2019).

A common approach for mapping LRS reads is seeding and chaining, where an approximate alignment is formed based on a subset (chain) of exact matches between the sequence and the genome. The exact matches may be found using various data structures including variable-length matches using a BWT-FM index or suffix arrays (Chaisson and Tesler, 2012), and minimizer based indexing (Li, 2018). The chaining algorithm used by BLASR uses a linear cost gap function for sparse-dynamic programming (SDP) (Baker and Giancarlo, 2002). While the chaining algorithm is efficient, it has long been known that linear-cost gap functions do not accurately reflect biological variation (Fitch and Smith, 1983), and has been shown to decrease sensitivity for detecting SV from LRS alignments (Sedlazeck *et al.*, 2018).

Both the minimap2 and ngmlr aligners use gap penalties that are a convex function of gap length, and are demonstrated to be quite effective for mapping LRS reads across SV with alignment gaps that reflect biological variation. In minimap2, a heuristic algorithm is used for chaining, while ngmlr adopts a Smith-Waterman-like dynamic programming algorithm. An exact solution to sparse dynamic programming with a convex gap (CG-SDP) function exists (Eppstein *et al.*, 1992b), and is slightly less efficient than linear-cost SDP. However, as presented, the algorithm requires asynchronous processing and has never been implemented for sequence alignment in computational biology. Furthermore, the algorithm does not take into account genome arrangements such as inversions.

An additional challenge for alignment enabled by LRS is discovering variation from *de novo* assembly. Efficient LRS assembly algorithms enable assembly of human genomes with contiguity on part of the initial release of the human genome (Shafin *et al.*, 2020; Kolmogorov, 2019). When combined with single-nucleotide polymorphisms phased using trio or long-range sequencing information such as Hi-C and strand-seq, LRS may be assembled into phased diploid genomes (Koren, 2018; Porubsky *et al.*, 2019). The breakpoints of sequence variation may be detected directly from assembly alignments, as opposed to read based variant detection which requires the consensus sequence of noisy breakpoint sequences that may have reference bias. This benefit comes at the expense of requiring accurate alignment of sequences tens of millions of bases long that may harbor rearrangements and duplications. The MashMap method can rapidly detect sequence homology in assemblies with low memory requirements (Jain *et al.*, 2018), however the pairwise alignments are not detailed enough to detect variants less than a few hundred bases, which accounts for the majority of variation between human genomes. It is also possible to align long contigs using minimap2, which can produce output that captures all scales of variation from SNPs to large SV, however the resulting alignments tend to be split at structural variation and can be mismapped in repetitive regions.

To explore the application of the exact solution of seed chaining sparse dynamic programming with a convex gap function to read and assembly alignment, we developed an alignment method lra. We have simplified the implementation of the CG-SDP algorithm (Eppstein et al., 1992b), and extended to allow for inversions and translocations.

## 2 Results

We compared alignment metrics and variant discovery on data from HG002 including three sequencing datasets: PacBio consensus reads (HiFi), PacBio single-pass reads (CLR), and ONT reads, as well as on contigs from a haplotype-resolved de novo assembly (Koren, 2018). All data were mapped to build GRCh37 to use curated variants for accuracy analysis (Zook *et al.*, 2020). The PacBio HiFi data are characterized by high-accuracy reads (>99%) with an average length of 19kb, compared to the CLR reads that have an accuracy around 80% and an average read length of 21kb. The ONT data: 88% accuracy and 1.2kb average read length. All read datasets are above 40 × coverage of a human genome, which are sufficient for human sequencing studies.

We compared read alignments to minimap2 and ngmlr, two commonly used alignment methods for LRS data. When computing SAM formatted alignments with minimap2, the lra and minimap2 runtimes are comparable; lra is faster for aligning HiFi reads, while minimap2 is faster for CLR and ONT data, although all are within a factor of two (Table 1). The number of bases aligned by lra and minimap2 are within 0.03-7%. While the performance metrics for runtime and number of aligned bases are worse for ngmlr, it is important to consider this method for downstream analysis because the potential for sensitivity for SV detection may be more important than runtime. A mapping quality is assigned based on the number of mismatches of secondary alignments close to the same length of the primary alignment. Simulated data was used to measure how mapping accuracy tracks with mapping quality. We compared the mapping accuracy of lra and minimap2 on simulated HiFi, CLR and ONT data of different read lengths: 5kb, 10kb, 20kb and 50kb. Simulated reads were mapped to genome GRCh38. A read is considered as correctly mapping if its longest alignment has ? 30% overlap with its true interval. The mapping accuracy is measured by *NC/TN*, where *NC* is number of correctly mapped reads and TN is total number of mapped reads (Figure 1). Reads were simulated using a method, alchemy2 included with the lra source that builds an error model from a bam file of aligned reads. While alignment accuracy was up to 0.5% lower than that of minimap2 in repetitive regions (indicated by lower mapping quality), downstream analysis was not affected.

**Fig. 1.**
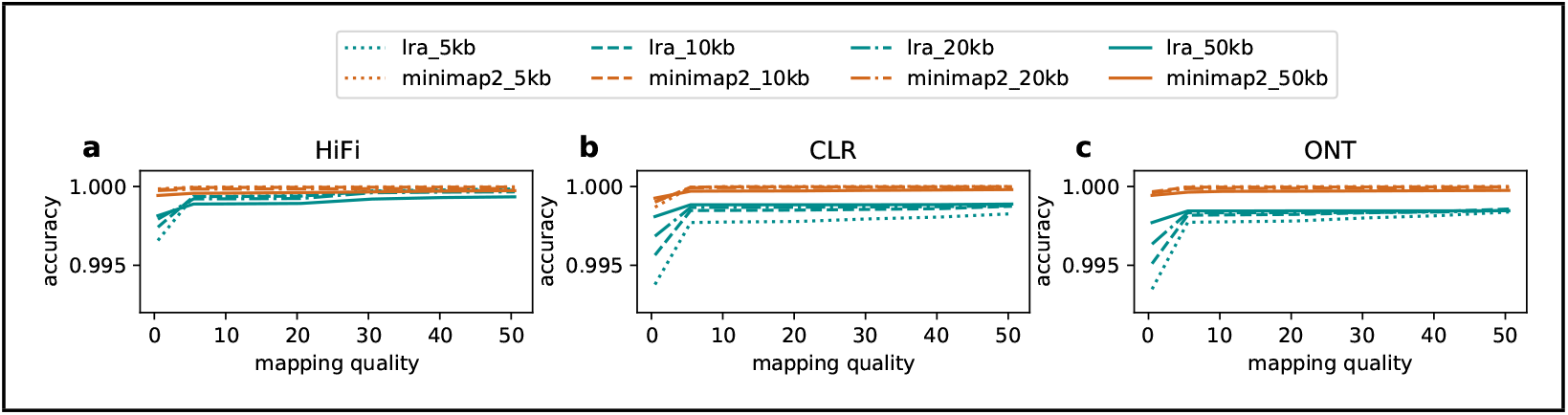
Mapping accuracy of lra and minimap2 on simulated HiFi, CLR and ONT reads of different lengths: 5kb, 10kb, 20kb 50kb. Simulated reads were mapped to genome GRCh38. A read is considered as correctly mapping if its longest alignment has ≥ 30% overlap with its true interval.

**Fig. 2.**
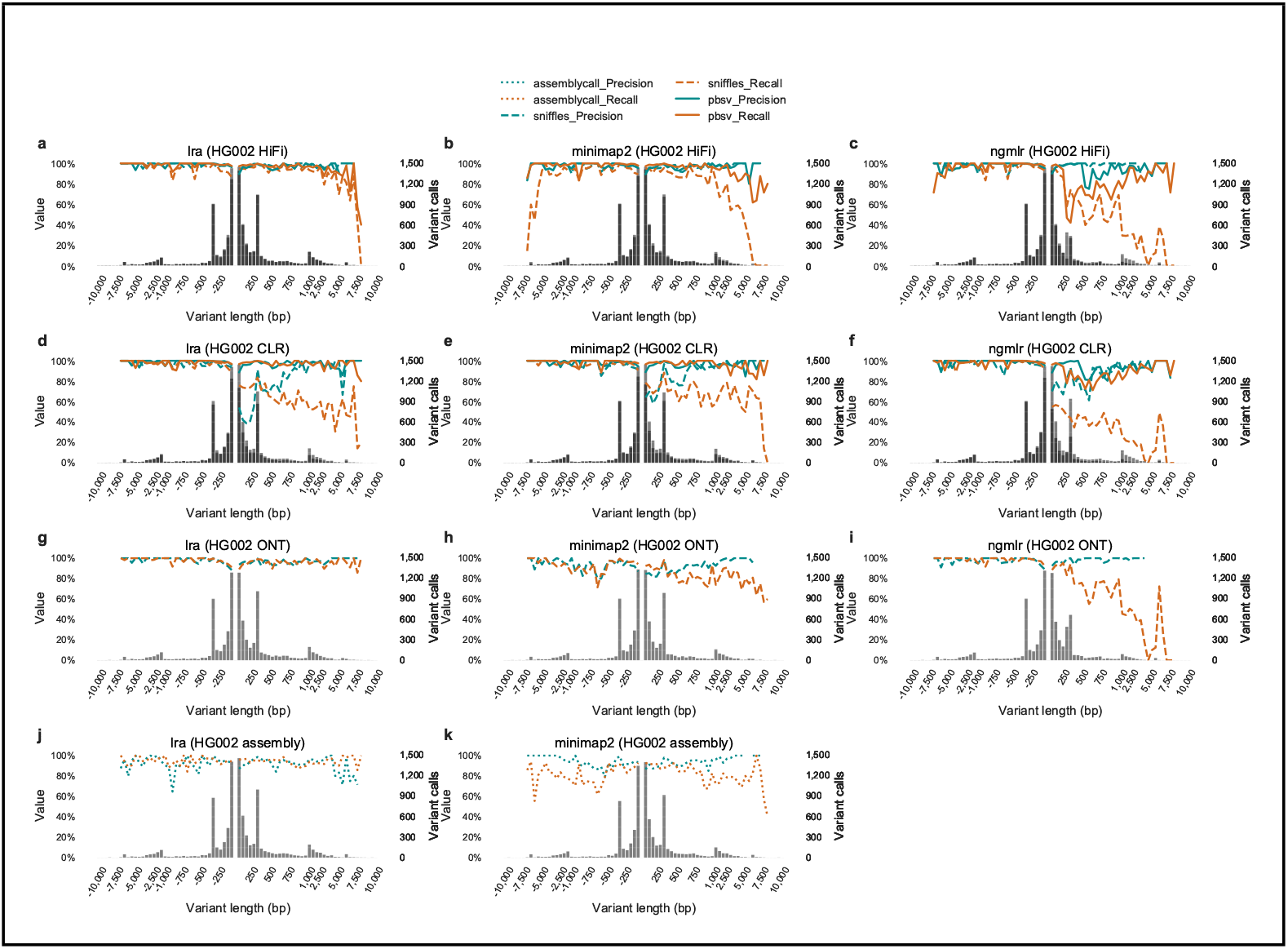
The precision and recall of SV call sets from HG002 HiFi, CLR, ONT, and assembly-based callsets, measured by Truvari using Genome in a Bottle benchmark SV callsets.

**Table 1.**
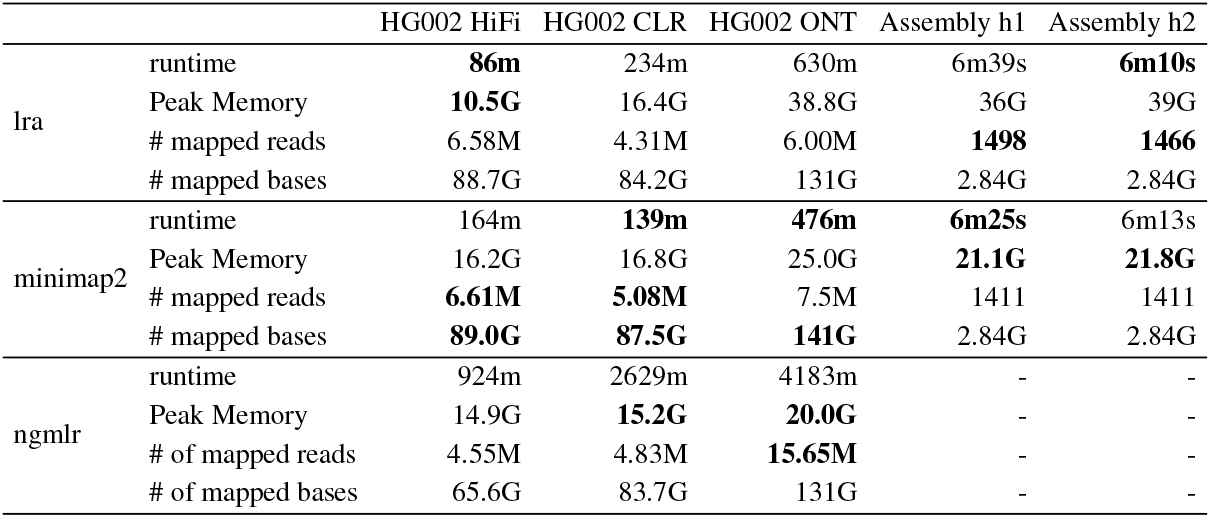
Performance of alignment methods. Each dataset was aligned to GRCh37 by all methods with 16 threads, with - indicating a software crash. The optimal values in each class are given in bold. The HiFi, ONT, and CLR data are 40, 42, and 79 fold coverage, respectively. The assembly is a haplotype-resolved de novo assembly of HG002 using HiFi reads, with haplotype N50 values of 16.1Mb and 18.0Mb.

|  |  | HG002 HiFi | HG002 CLR | HG002 ONT | Assembly h1 | Assembly h2 |
| --- | --- | --- | --- | --- | --- | --- |
| lra | runtime | **86m** | 234m | 630m | 6m39s | **6m10s** |
|  | Peak Memory | **10.5G** | 16.4G | 38.8G | 36G | 39G |
|  | # mapped reads | 6.58M | 4.31M | 6.00M | **1498** | **1466** |
|  | # mapped bases | 88.7G | 84.2G | 131G | 2.84G | 2.84G |
| minimap2 | runtime | 164m | **139m** | **476m** | **6m25s** | 6m13s |
|  | Peak Memory | 16.2G | 16.8G | 25.0G | **21.1G** | **21.8G** |
|  | # mapped reads | **6.61M** | **5.08M** | 7.5M | 1411 | 1411 |
|  | # mapped bases | **89.0G** | **87.5G** | **141G** | 2.84G | 2.84G |
| ngmlr | runtime | 924m | 2629m | 4183m | - | - |
|  | Peak Memory | 14.9G | **15.2G** | **20.0G** | - | - |
|  | # of mapped reads | 4.55M | 4.83M | **15.65M** | - | - |
|  | # of mapped bases | 65.6G | 83.7G | 131G | - | - |

**Table 2.** Classification of inversions detected in HG002 using PacBio and ONT reads. TP, true positive using manual curation. Dup, inverted duplication misclassified as an inversion on both haplotypes, NA not possible to manually curate, FP - no signature of inversion in read nor assembled haplotype dot-plots.

| Dataset | Aligner | TP | Dup | NA | FP |
| --- | --- | --- | --- | --- | --- |
| CCS | lra | 37 | 50 | 1 | 6 |
| CCS | mm2 | 24 | 1 | 3 | 2 |
| CCS | ngmlr | 70 | 31 | 2 | 6 |
| CLR | lra | 34 | 6 | 2 | 6 |
| CLR | mm2 | 36 | 7 | 8 | 3 |
| CLR | ngmlr | 78 | 29 | 4 | 17 |
| ONT | lra | 19 | 2 | 1 | 2 |
| ONT | mm2 | 30 | 7 | 1 | 4 |
| ONT | ngmlr | 60 | 20 | 22 | 0 |

A common application of LRS in human genetics is detecting structural variation (Chaisson et al., 2019; Sedlazeck et al., 2018). We evaluated SV detection from mapped reads across combinations of technologies, aligners, and SV calling algorithms. Reads were aligned with lra, pbmm2/minimap2, and ngmlr, and variants were detected with pbsv and Sniffles (Wenger et al., 2019; Sedlazeck et al., 2018). The pbmm2 method encapsulates minimap2 with technology and SV discovery specific parameters. The pbsv method did not run using ONT reads, and so pbsv analysis of ONT data is omitted. The majority of SV are under 500 bases (Chaisson *et al.*, 2019) and are spanned by LRS alignments. Consequently, variant calls with similar representations of gaps should have converging callsets, which may be confirmed by comparing SV calls from different combinations of algorithms. The most consistent variant calling was found using pbsv on both types of PacBio using lra and pbmm2 alignments. These callsets ranged from 20,574-21,754 SVs, with a difference of 1.4-1.7% between alignment algorithms and 1.5-4.6% between data types (Table 3). Compared to the pbsv calls, there was greater variation between callsets generated using Sniffles across both data types and alignment algorithms, with a range of 15,062-24,067 SVs per callset. There was less variation between the size of callsets of different data types and the same alignment algorithm (average difference of 2,767 call differences per callset), compared to different alignment algorithms applied to the same data (an average difference of 6,412 per callset).

**Table 3.** Structural variant call sets. Summary of SV callsets from different aligners (lra/pbmm2/ngmlr)

|  |  |  | lra |  |  | pbmm2 |  |  | ngmlr |  |  |
| --- | --- | --- | --- | --- | --- | --- | --- | --- | --- | --- | --- |
|  |  |  | Number of calls | Average call size | Total size | Number of calls | Average call size | Total size | Number of calls | Average call size | Total size |
| pbsv | CLR | INS | 12328 | 509 | 6.27M | 12590 | 524.23 | 6.60M | 10123 | 672 | 6.81M |
|  |  | DEL | 8426 | 581 | 4.90M | 8520 | 518.39 | 4.42M | 9172 | 513 | 4.71M |
|  | HiFi | INS | 12655 | 489 | 6.19M | 12828 | 523.53 | 6.72M | 11508 | 398 | 4.59M |
|  |  | DEL | 8784 | 583 | 5.12M | 8926 | 603.51 | 5.39M | 9960 | 520 | 5.18M |
| Sniffles | CLR | INS | 13247 | 499 | 6.60M | 11186 | 397.337 | 4.44 | 6638 | 308 | 2.04M |
|  |  | DEL | 10820 | 678 | 7.33M | 9264 | 685.866 | 6.35 | 8424 | 1054 | 8.87M |
|  | HiFi | INS | 11995 | 604 | 7.23M | 10779 | 380.117 | 4.09 | 8064 | 262 | 2.11M |
|  |  | DEL | 8797 | 802 | 7.05M | 7865 | 458.806 | 3.6 | 7685 | 913 | 7.01M |
|  | ONT | INS | 12299 | 714 | 8.78M | 12299 | 415.157 | 4.97M | 8800 | 336 | 2.95M |
|  |  | DEL | 10005 | 688 | 6.88M | 9317 | 684.443 | 6.37 | 8315 | 822 | 6.83M |

A greater call count may reflect more sensitive detection of SV, or simply fragmentation of variants. To assess the accuracy of variant callsets, we compared callsets against the GIAB Tier 1 calls (Zook *et al.*, 2020) using Truvari (https://github.com/spiralgenetics/truvari). Thelraand pbmm2 based callsets outperform ngmlr based calls in precision and recall on all data types (Table 4). While the F1 scores are effectively equivalent between lra and pbmm2 based variant calls on HiFi (0.972 vs 0.968) and CLR (0.969 vs 0.967) using pbsv, there is a substantial improvement on calls on ONT data made by Sniffles (0.950 vs 0.910). This indicates that the greater call count on ONT data includes is at least partially attributed to increasing recall, particularly in larger (>500 base) insertions, without affecting precision on the high-quality regions that were ascertained.

**Table 4.** Truvari classification of variant calls. Truvari comparisons between lra and pbmm2/minimap2 using the Genome in a Bottle benchmark SV set. Optimal results in each category are shown in bold. TP-base means true positive calls in the benchmark SV curation set, while TP-call means true positive calls in the SV set from each aligner. False positive means the number of non-matching calls from the SV set from each aligner. False negative means the number of non-matching calls from the SV curation set. False negative variants that show underlying read support are given by s/FN.

|  | pbsv |  |  |  |  |  | Sniffles |  |  |  |  |  |  |  |  |
| --- | --- | --- | --- | --- | --- | --- | --- | --- | --- | --- | --- | --- | --- | --- | --- |
|  | HiFi |  |  | CLR |  |  | HiFi |  |  | CLR |  |  | ONT |  |  |
|  | lra | pbmm2 | ngmlr | lra | pbmm2 | ngmlr | lra | pbmm2 | ngmlr | lra | pbmm2 | ngmlr | lra | mm2 | ngmlr |
| TP base | **9425** | 9408 | 8433 | 9397 | **9414** | 9076 | **9012** | 8700 | 8015 | 7852 | **8082** | 6533 | **9123** | 8887 | 8471 |
| TP call | **9459** | 9449 | 8500 | 9484 | **9507** | 9208 | **9034** | 8713 | 8028 | 7942 | **8147** | 6567 | **9151** | 8902 | 8488 |
| FP | **331** | 390 | 396 | **358** | 418 | 428 | **281** | 357 | 800 | 2705 | 1492 | **1298** | **445** | 940 | 815 |
| FN | **216** | 233 | 1208 | 244 | **227** | 565 | **629** | 941 | 1626 | 1789 | **1559** | 3108 | **518** | 754 | 1170 |
| s/FN | 124 | 93 | 176 | 167 | 151 | 268 | 516 | 598 | 573 | 1706 | 1423 | 2387 | 439 | 464 | 410 |
| precision | **0.967** | 0.96 | 0.955 | **0.964** | 0.958 | 0.956 | **0.97** | 0.961 | 0.909 | 0.746 | **0.845** | 0.835 | **0.954** | 0.904 | 0.912 |
| recall | **0.978** | 0.976 | 0.875 | 0.975 | **0.976** | 0.941 | **0.935** | 0.902 | 0.831 | 0.814 | **0.838** | 0.678 | **0.946** | 0.922 | 0.879 |
| F1 score | **0.972** | 0.968 | 0.913 | **0.969** | 0.967 | 0.948 | **0.952** | 0.931 | 0.869 | 0.779 | **0.842** | 0.748 | **0.95** | 0.91 | 0.895 |

While the F1 scores are nearly equivalent on PacBio data, the combination of algorithms may contribute to a more complete evaluation of an LRS genome. The lra and pbmm2 contribute 67 and 59 unique calls, respectively, that were annotated as true positives in the HiFi/pbsv call sets, and 83/112 unique calls from the CLR/pbsv call sets. The average length of the uniquely called true positive SV is between 1.3-4.4kb, highlighting the challenge of detecting longer SV from read based alignments. To further assess the potential for callsets to be more comprehensive, benchmark variants annotated as false-negatives were compared to variants discovered directly from read alignments. Each call where at least 20% of the reads overlapping the call had an SV of the same type with a length between 0.6-1.4 × the length of the call were annotated as a supported false-negative (s/FN). Between 14.5%-68.4% of annotated false negatives from pbsv variant callsets, and 35.0%-95.4% of false-negative calls in Sniffles callsets were considered s/FN. Furthermore, there were 44% more supported false-negative calls in the PacBio HiFi lra/pbsv callsets compared to the pbmm2/pbsv callsets, and on average 79% more s/FN calls from lra and minimap2 alignments in Sniffles callsets, an expected outcome of tuning a variant discovery algorithm to input generated by a particular alignment algorithm.

A comparison of inversion calls between algorithms indicates this class of variation is a remaining challenge for SV discovery. Inversion calls for ngmlr alignments were directly obtained from the vcf file generated by Sniffles. For lra and minimap2, inversions calls were first obtained from the PAF output, since a tag indicating inversion is given in the PAF and the same calls from different reads alignments were merged. We filtered calls that overlapped the centromere, assembly gaps, or sequence coverage (PacBio CLR lra alignments) greater than twice the average coverage. Each resulting callset was manually curated using dot-plots of the genome assemblies or HiFi reads against the reference, and using samplot (Belyeu *et al.*, 2020). Calls were classified as true positives if at least one haplotype or an exemplary HiFi read clearly showed the inversion or it was indicated from the samplot alignments, a duplication if the dot-plot signature was a fixed inverted duplication or inverted transposition, false-positive if both haplotypes did not show variation or an inverted duplication structure, and NA if it was not clear, commonly in pericentric regions. False calls by any method may be easily filtered by comparing alignments using the corresponding inversion sequence in forward an reverse orientation across multiple reads. Across all data types, ngmlr based alignments discovered the most inversions annotated as true-positives, with calls ranging from 77-91,152 bases (Table 2). On average, 69 inversions were detected using ngmlr alignments, versus 30 for both lra and minimap2 alignments, indicating that additional development may be required to accurately detect rearranged DNA with minimal computational burden.

When sequencing depth is sufficiently high coverage, comprehensive haplotype-resolved de novo assemblies can be used to detect variation instead of read alignments. Both lra and minimap2 were used to align contigs of a haplotype-resolved de novo assembly of HG002 constructed + SV callers (pbsv/Sniffles).by the trio-canu assembler (Wenger *et al.*, 2019). The haplotype assemblies resolve 2.96Gb and 3.04Gb, with N50 values 16.1 and 18.0 Mb. Both alignment methods have similar runtimes to align *de novo* assembly contigs (< 7 min, 16 threads), although lra requires roughly twice the memory. It was not possible to use ngmlr to align contigs. Variants were detected from the minimap2 alignments using paftools.js. To detect variants using lra, alignments were first filtered by removing short alignments spanning less than 200kb, and calling SV directly from alignments (e.g. CIGAR strings), selecting calls only from the longest contig when multiple contigs overlap. Calls from different haplotypes that overlap at least 30% were determined as homozygous calls and the rest were classified as heterozygous calls. Additionally, calls in centromere regions were removed for both minimap2/paftools.js and lra callsets. This generated 25,708 calls by lra call sets, and 24,149 by minimap2/paftools.js. Truvari analysis of the calls gives an F1 score of 0.9382 by lra calls, and 0.8989 for pbmm2 calls (Table 5).

**Table 5.** Assembly based calls. Calls from lra and minimap2 assembly alignments are classified as homozygous and heterozygous by comparing calls from two haplotypes. Truvari comparison results between these two callsets are shown.

| HG002 assembly |  |  |
| --- | --- | --- |
|  | lra + pruning | minimap2 + paftools.js |
| Insertion, hom | 5415 | 5784 |
| Insertion, het | 9320 | 8582 |
| Deletion, hom | 2864 | 3137 |
| Deletion, het | 7325 | 6047 |
| TP base | **9118** | 8479 |
| TP call | **9319** | 8512 |
| FP | **693** | 747 |
| FN | **523** | 1162 |
| precision | **0.9307** | 0.9193 |
| recall | **0.9457** | 0.8794 |
| F1 score | **0.9382** | 0.8989 |

**Table 6.** Read based support of assembly calls. The two haplotype assemblies are considered separately to avoid complications of merging into a diploid callset. Calls are produced by lra and minimap2 (mm2), with only deletion and insertion SV classes considered. The total calls are the original calls produced by each method, and filtered are calls excluding centromeres and 50kb of flanking sequence. The supported SV have at least four reads supporting the call from either lra or minimap2 alignments.

| Assembly | SV type | total | filtered | supported (fraction) |
| --- | --- | --- | --- | --- |
| Haplotype 1, lra | DEL | 6151 | 5988 | 5900 (0.985) |
| Haplotype 1, mm2 | DEL | 5808 | 5804 | 5782 (0.996) |
| Haplotype 2, lra | DEL | 6074 | 6029 | 5944 (0.986) |
| Haplotype 2, mm2 | DEL | 5680 | 5671 | 5645 (0.995) |
| Haplotype 1, lra | INS | 9760 | 9589 | 9522 (0.993) |
| Haplotype 1, mm2 | INS | 9640 | 9634 | 9624 (0.999) |
| Haplotype 2, lra | INS | 9546 | 9496 | 9435 (0.994) |
| Haplotype 2, mm2 | INS | 9394 | 9383 | 9368 (0.998) |

The high-confidence callset used in Truvari analysis contains fewer than 10,000 SVs, less than half what are expected to be found in a human genome. To gauge the specificity of assembly-based calls outside this set, SV contained in read alignments were compared to the SV discovered from assemblies. An SV detected from an individual read supported an assembly based SV if the call was the same type (e.g. insertion or deletion), was within 1kb, and the ratio of SV lengths was between 0.5 and 2. When using PacBio CLR lra-aligned reads, the majority (99.7%) of SV annotated by Truvari as true-positives had at least four supporting reads. The overall support is defined from the union of support from both lra and minimap2 alignments in order to mitigate any alignment bias. Using this approach, 98.7% of lra deletion SV, and 99.7% of minimap2 deletion SV are supported by reads, and 99.6% / 99.9% of insertion SV from lra/minimap2, are supported by reads 6. This is greater than the precision measured by Truvari on assembly-based calls: 0.9332 for lra, and 0.9295 for minimap2, indicating an under estimates of the callset precision. When inspecting the SV calls that are not supported by read alignments, many are due to differential placement of gaps causing disagreement of SV length between the read and assembly-based calls.

## 3 Methods

The alignment of reads and contigs to a reference are generally defined by the maximally scoring local alignment of a query *q* to a set of target sequences collectively referred to as a target *t* with a match bonus and penalties for mismatch, gaps, and inversions/translocation. lra uses a heuristic to find an approximate local alignment employing the commonly used seeding, chaining, and refinement approach that has been applied to all scales of alignment from short-read, long-read, and whole-genome alignment (Kent *et al.*, 2003; Sedlazeck *et al.*, 2013; Chaisson and Tesler, 2012). Each alignment proceeds in four broad steps: seed sequence matching, clustering, chaining, and finally alignment refinement.

Many recent advances have been made in sequence mapping using a subsampled index on a reference using minimizers or locally sensitive hashing (Roberts *et al.*, 2004; Jain *et al.*, 2017). A minimizer index is parameterized by a k-mer size *k* and window size *w*, and indexes the position of the lexicographically least canonical k-mer in every sequence of length w across the genome. We develop a variant on the approach of minimizers that uses adaptive thresholds to limit the total number of positions sampled in unique regions of a genome, and increase the sampling of positions near paralog-specific variants that distinguish repetitive regions.

Optimal sets of matches between the query and reference are selected in two phases using CG-SDP. Given a set of fragments Φ = {*α*_1_,…, *α_n_* }, each fragment is defined by a start point and an end point on a Cartesian plane and a weight: 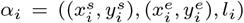. The basic CG-SDP formulation is to define a chain of fragments *C* ⊂ [1,…, *n*] that maximizes

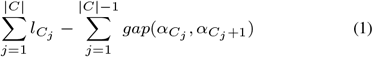

such that *α*_*C*_*j*+1__ is above and to the right of *α*_*C*_*j*__, and *gap* is a convex function. An algorithm was presented for chaining using a convex gap cost model (Eppstein *et al.*, 1992b), however there are no alignment methods that implement this approach. This method has been implemented in lra and extended to allow for inversions in the optimal chain. Additionally, the original description of the algorithm requires asynchronous processing, which we have updated to use standard serial computation. The details of determining minimizers and the chaining algorithm are given below.

### 3.1 Building a Minimizer Index and Sequence Matching

We add three additional parameters to generate a minimzer index: *F^M^*, *N^M^* and *W^G^* that limit the density of minimizers that are selected. An initial set of minimizers is determined in the standard approach, with minimizer k-mer parameter *k* and window *w* (Roberts *et al.*, 2004). Next, minimizers of multiplicity larger than *F^M^* are removed. Then the reference is partitioned into intervals of length *W^G^*, and the remaining minimizers starting in each interval are selected in order of their multiplicity in the genome until the first *N^M^* minimizers are obtained. lra uses slightly different parameters settings for aligning reads from different sequencing technologies to genome. When indexing the genome for aligning HiFi reads, 867M minimizers from total 1015M were left after filtering by frequency threshold *F^M^*. In total, 117M minimizers were selected after the final filtering based on *N^M^* and *W^G^*. All minimizers from a query sequence are matched against the filtered set of minimizers from the reference. The result of the sequence matching is a set of anchors *A* = {*β*_1_,…, *β_n_*}, where *β_i_* is a tuple (*x_i_*, *y_j_, k*), where *q* [*x_i_*, *x*_*i*+1_,…, *x*_*i*+*k*−1_] of the query matches *t* [*y_j_*, *y*_*j*+1_,…, *y*_*j*+*k*-1_] of the target.

An additional local minimizer index is built on the query and target, parameterized by standard minimizer parameters *k^l^* and *w^l^,* and a tiling length *W^l^.* A separate minimizer index is stored for sequences of length *W^l^* tiling across both the query and the target, with *k^l^* < *k* and *w^l^* < *w*. The local minimizer index is used to speed the alignment of the query and target once the rough alignment is detected.

### 3.2 Clustering

Although CG-SDP can be applied to all anchors A, for efficiency a greedy approach is used to cluster anchors that would likely be together on an optimal chain, and CG-SDP is applied to clusters. Two clustering steps are performed, rough and fine clustering, where rough clusters contain matches in the same target region, and fine clusters divide rough clusters into non-overlapping sets of anchors that are not likely to contain large insertion, deletion, or rearrangement variants.

#### 3.2.1 Rough Clustering

Rough clustering partitions anchors into clusters representing approximate intervals on the query and target that are aligned (Figure3a). Denoting the forward diagonal of each anchor *β_i_* as *f_i_* = *y_i_* – *x_i_*, and the reverse diagonal *r_i_* = *x_i_* + *y_i_*, a sorted anchor order *O* = [*o*_1_,…, *o_n_*] is defined by ordering anchors by forward diagonal and then x coordinate. A reverse sorted order 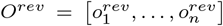 is similarly defined sorting on reverse diagonal and x coordinate. This will be used to detect alignments on the reverse strand, but because the operations are the same as on the forward strand, only subsequent steps using the forward sorted order are given. The set of rough clusters is defined by partitioning O into non-overlapping intervals such that every anchor indexed in an interval has a diagonal within *D^R^* of the first anchor indexed in the interval. Intervals are greedily assigned with first interval starting at the first index in O, and subsequent intervals starting on the first anchor beyond *D^R^* of the previous interval. Intervals with few elements (defined by a *minClusterSize* parameter) are discarded, and the rough clusters *R* = {*R*_1_,…, *R_NR_*} are defined from the set of anchors included in each interval.

#### 3.2.2 Fine clustering

In this step, each rough cluster is processed independently by dividing into non overlapping fine clusters, where each fine cluster consists of anchors on a close diagonal *D^F^*, where *D^F^* < *D^R^*, with endpoints that do not overlap. The first step of CG-SDP will be applied to chain the fine clusters and find an approximate alignment between *q* and *t*. Each fine cluster *C_j_* is defined by all of the anchors contained in the cluster, and endpoints 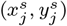 and 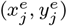, where 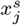, 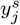 are the minimum *x, y* coordinates of the starting points of all the anchors in *C_j_*, and 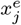, 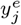 are the maximum *x, y* coordinates of the ending points of all the anchors in *C_j_*. To define the fine clusters, anchors in each rough cluster are first sorted by Cartesian coordinate. Within each rough cluster, an anchor is defined as unique when the k-mer of the match is not repeated in the cluster. Fine clusters are initialized as runs of unique anchors in the Cartesian ordering that are on a close diagonal, and the distance between the end of one anchor and the start of the next is small (Figure 3b). Every pair of fine clusters *C_j_* and *C_k_* are merged if their endpoints have diagonal differences smaller than *D^F^* and are within Cartesian distance *G^dist^* (Figure 3c), and all non-unique anchors within the trapezoid defined by
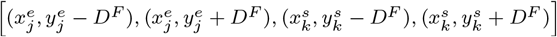 are included into the merged fine cluster (Figure 3d, 3e). The remaining non-unique anchors that are not added into the fine cluster are discarded.

### 3.3 Cluster Splitting and Chaining

Each fine cluster *C_j_* ∈ *C* defines a super-fragment *F_j_* that unlike the minimizer fragments which have starting and endpoints along the same diagonal, the endpoints of *C_j_* and may not be along a diagonal. The set of all such fragments is *F*, and an optimal chain of fragments will be defined using CG-SDP. However, considering the rectangles defined by the boundaries of each fragment, the CG-SDP algorithm only selects fragments with non-overlapping rectangles in the optimal chain. Due to the repetitive nature of genomes, this may result in erroneously skipped fragments (Figure 3). To account for this, fine clusters are split at overlapping boundaries (Figure 3g). The start and end coordinates of all fine clusters are stored in a set that is queried to find boundary points appearing in the range of each fine cluster. The coordinates of each split cluster are set according to the first/last anchors appearing after/before the boundary point. An optimal chain of fragments *F^SDP^* ∈ *F* with corresponding clusters *C^SDP^* are then found using CG-SDP on F.

### 3.4 Cluster Refinement

The optimal chain of super-fragments obtained from section 3.3 contains the anchors from which the optimal alignment will be defined. A pairwise alignment may be created using dynamic programming on the substrings between the minimizer matches, however for high-error rate reads matches are sparse and the length of substrings may be too large to efficiently compute, or have too large of a diagonal gap to use banded alignment. A second anchoring step using the local minimizer index is used to detect shorter and more dense anchors. The local minimizer index contains 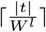 and 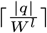 separate minimizer indexes for the target and query sequences that index each tiling substring of length *W^l^*, accounting for the relative positions of the substrings in each sequence. Every pair of substrings from q and t that have an anchor in *C^OPT^* are compared using their local minimizer indexes to generate a resulting set of anchors *A^local^*. To reduce runtime of CG-SDP on *A^local^*, anchors that are adjacent in Cartesian sorted order and on the same diagonal are merged. The length of any merged anchor is the difference from the last endpoint to the first starting point. The resulting merged fragments are chained using CG-SDP, as described in section 3.6, and the anchors on this chain are denoted as *A^local-opt^*

### 3.5 Alignment Refinement

Banded alignment is used to create a pairwise alignment between anchors in *A^local-opt^*. We assume that large gaps between anchors may be modeled using an affine alignment that allows a single large penalty-free gap. Given two sequences *q^local^* and *t^local^*, a match matrix *M*, a gap penalty *δ*, and an alignment band *B,* assume that |*q^local^*| < |*t^local^*| – *B*. A lower-score matrix *S^lower^* is calculated for a typical banded alignment band B and gap penalty *δ*, e.g 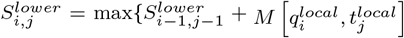, 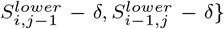 if |*i* – *j*| ≤ *B*, –∞ otherwise. Next, an upper-diagonal matrix 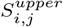 is calculated that allows for a single transition from the lower matrix with banded alignment. Denoting the length of the *q^local^* and *t^local^* as *l^q^* and *l^t^*:

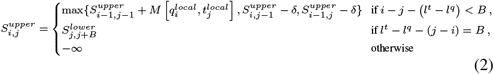

**Fig. 3.**
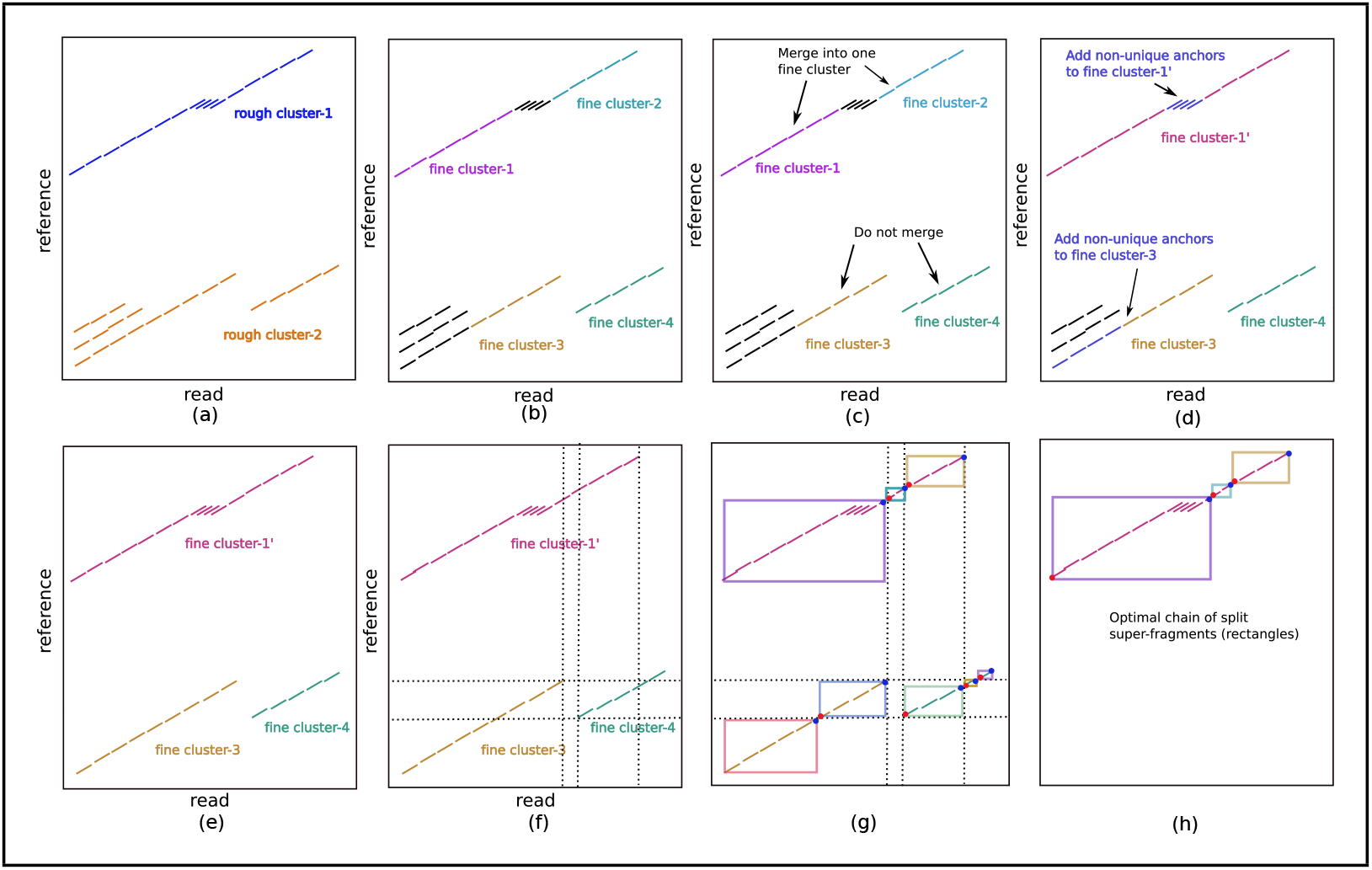
Example of clustering anchors prior to CG-SDP. a, Two rough clusters (blue and orange), which are far from each other on the reference. b, The initial four fine clusters defined from the contiguous stretches of unique anchors. c, Fine cluster-1 and fine cluster-2 are merged because their diagonal difference is smaller than *D^F^* and projected distances between their endpoints is smaller than *G^dist^*. Fine cluster-3 and fine cluster-4 are not merged due to the large diagonal difference. d, Non-unique anchors in the trapezoid between fine cluster-1 and fine cluster-2 are added to the merged fine cluster-1, along with non-unique anchors in the trapezoid defined by the start of rough cluster-2 and the start of fine cluster-3. e, Three fine clusters are obtained after rough clustering and fine clustering. f-h, Splitting of overlapping fine-clusters: f, overlap of clusters. g, Boundaries of split clusters defined by a start (red dot) and an end (blue dot). h, the optimal chain of split super-fragments.

### 3.6 Problem Statement of the Chaining

We define a set of fragments Φ = {*α*_1_,…,*α_n_*}. Each fragment *α_i_* is associated with a lower left start point 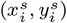 and upper right end point 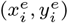, and a score *l_i_*. For minimizer fragments, the upper right endpoints are a fixed distance from the lower points, e.g. 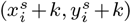. For super fragments defined by fine clusters, the endpoints and starting points may not be on the same diagonal. The starting point of a fragment *α* is denoted *s*(*α*), and the end *e*(*α*). A point (*x_i_*, *y_i_*) is above (*x_j_*, *y_j_*) if *x_i_* > *x_j_*, *y_i_* > *y_j_* (conversely (*x_j_*, *y_j_*) is below (*x_i_*, *y_i_*)). The chaining score is defined by Equation 1 where *gap*(*α_C_j__*, *α*_*C*_*j*+1__) is a convex function of 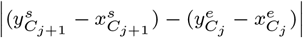, the difference between the forward diagonals of the endpoint of *α_C_j__* and the starting point of *α*_*C*_*j*+1__. The naive way to solve this problem takes *O*(*n*^2^) in time. By applying CG-SDP (Eppstein et al., 1992b), the runtime is *O*(*n* log(*n*)^2^). Below, the solution provided by Eppstein, Galil, and Giancarlo is described both with increase clarity from the original description, with a modification that enables calculation with synchronous computation. Finally, the method is extended to allow for inversions, similar to Brudno *et al.* (2003).

#### 3.6.1 SDP Algorithm with Convex Gap Cost Function - Defining Subproblems

For simplicity, assume both sequences are the same length l and that all points are in [0, *l*) (e.g. shifted by the minimum *x* and *y* coordinate), and are on a *l* × l grid. To speed the chaining algorithm, the search for the fragment that precedes another on an optimal chain is divided into multiple overlapping subproblems that may be solved independently and more efficiently than the naive scan, and the globally optimal score for each point is selected from each of the subproblems that overlap it. Each subproblem divides a block of *t^sub^* columns or rows of the search space into an *A*-part and *B*-part covering 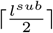 and 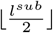 columns/rows respectively, where *A*-part contains all the endpoints and *B*-part contains all the startpoints in the corresponding columns/rows. When the size of a subproblem is only one column/row, the *A*-part of the corresponding subproblem is set to be empty. Each subproblem is described by the label *d* ∈ {*column, row*}, the starting and ending rows and columns of the subproblem *A.s, A.e, B.s, B.e*, and a data structure of values used to compute the optimal chain, *DATA*. The full set of subproblems *Sub*(*d, s, e*) are generated recursively as:

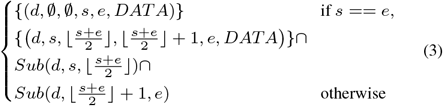

The more detailed pseudocode for this partitioning is given in the supplementary material (Supplementary Alogrithm S1). Figure 4 visualizes the column and row subproblem division of a simple case with six fragments.

Each subproblem is processed by finding the optimal endpoint from the *A*-part that precedes each starting point in corresponding *B*-part. For a starting point *p_i_* that is assigned to the *B*-parts of a set of column-based subproblems, the union of the *A*-parts of those subproblems form the entire plane to the left of the point. Similarly for a starting point assigned to the *B*-part of a set of row subproblems, the union of the *A*-parts of those subproblems form the plane below the point. After solving for the optimal preceding endpoint in all of the subproblems in which *p_i_* is contained, the one with maximum score among these endpoints is the global optimal, which represents the optimal chain from all endpoints below and to the left of *p_i_*.

Once the subproblems are defined, list elements of *DATA* are allocated and initialized. The following elements are associated with the *A*-part of a column/row subproblem:

- *D_I_*: The forward diagonals in increasing/decreasing order overlapped by endpoints in the *A*-part.
- *D_V_*: The optimal chaining score for diagonals overlapped by endpoints in *A*-part. *D_V_* [*s*] holds the optimal value of forward diagonal *D_I_* [*s*].
- *D_P_*: Index of point with optimal score along diagonal. *D_P_* [*s*] references the point with the best score of *D_V_*[*s*].

The following elements are associated with the *B*-part of a column/row subproblem:

- *E_I_*: The forward diagonals in increasing/decreasing order overlapped by starting points in the *B*-part.
- *E_V_*: Similar to Dy, the locally best chaining value.
- *E_P_*: Index of the forward diagonal in *D_I_* that leads to the best chaining value *E_V_*[*s*] for the forward diagonal *E_I_*[*s*].
- *E_B_*: A block structure used in the calculation of *E_V_*.
- *E_L_*: Index of the most recent value in *D_V_* used to compute values in *E_V_*.

Additionally, for each point, two lists hold references to each subproblem a point is contained in, one structure SA for points in *A*-parts of subproblems, and *SB* for points in *B*-parts of subproblems. To make the description of the method more concise, the operation *φ*(*D_I_, f*) and *φ*(*E_I_, f*) are defined to give the index of diagonal *f* in *D_I_* and *E_I_*, respectively.

#### 3.6.2 SDP Algorithm with Convex Gap Cost Function - Conquering Subproblems

The original description of CG-SDP, uses asynchronous processing during solving subproblems (Eppstein *et al.*, 1992b). Here we give an alternative description of the algorithm, and provide an approach to solve subproblems in a way that allows synchronous processing of points in Cartesian order for a more simple implementation. The framework for solving a subproblem is first described, and then an order of processing points is given to solve for optimal fragment chaining.

For any subproblem, the optimal chains between endpoints in the *A*-part to starting points in the *B*-part are found using the *D_I_, D_V_, D_P_, E_I_, E_V_, E_P_* arrays, variable *E_L_* and block list E_B_. Consider two points *p_a_* in *A*-part, and *p_b_* in *B*-part that are on diagonals *f_a_* and *f_b_*, respectively. An invariant for any subproblem is that if the optimal chain ending with the fragment that has endpoint pa has been solved and is referenced as *Score*(*p_a_*), the score of chaining *p_a_* with *p_b_* is *Score*(*p_a_*) + *gap*(|*f_b_* – *f_a_*|). Because the gap cost only depends on diagonal and not the coordinates of a point, all points in *A*-part on the same diagonal *f_a_* will have the same cost to chain with *p_b_*. More generally, the score to chain any starting point in *B*-part to an ending point in *A*-part is equal to the score of chaining a diagonal in *B* to a diagonal in *A*. The score of an optimal chain up to point *p_b_* in *B*-part, *E_V_* [*j*], where *j* = *φ*(*E_I_, f_b_*), is found for column-based problems as:

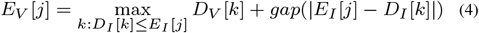

For row-based subproblems:

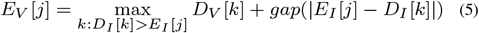

The naive approach to solve for all values of *E_V_* scales quadratically as *O*(|*E_V_* | | *D_V_*|). A problem of this form:

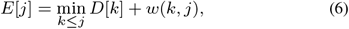

where *w* is a convex function and *D*[*k*] is a linear transformation of *E*[*k*], was solved in *O* (|*D*| log |*E*|) time using the auxiliary block data structure *E_B_* (Galil and Giancarlo, 1989; Gusfield, 1997). In this solution, iteration is performed with one pass over the *D* array. After each iteration *i*, the invariant holds: each element in *E* has been set to reference which element of the prefix *D*[1,…, *n*] gives the minimum value for Equation (6). Efficiency is gained by not updating E explicitly but by storing indexes in the data structure *E_B_* that may be updated in log(|*E*|) time by applying a function *Update* (*D* [*i*], *E_B_*) on each iteration. The data structure *E_B_* has an operation that can reconstruct *E*, *E*[*k*] = Ω(*E_B_, k*). The details of *E_B_* and *Update* may be derived from (Gusfield, 1997). In the application to CG-SDP, *D* is replaced by *D_V_* and *E* by *E_V_*, and Equation (4) and Equation (5) solve for the optimal chains between diagonals from an *A*-part of a subproblem to diagonals in the *B*-part of the column and row subproblem respectively. An important detail in our implementation is that in column-based problems an *Update* operation only affects references in *E_V_* that are on a greater or equal diagonal than the current element in *D_V_*, and for row-based problems elements in *E_V_* are only affected for diagonals that are less than the current element in *D_V_*. However, it is not possible to apply Equation (4) and Equation (5) directly. Because points are processed in Cartesian order, but *D_V_* in forward diagonal order, values of *D_V_* are not solved in increasing order and not all values of *D_V_* are guaranteed to be solved when values of *E_V_* are computed. In (Eppstein *et al.*, 1992b) this is accounted for by asynchronous computation. Below, an approach is described to solve with a standard model of computation by calling *Update*(*D_V_* [*i*], *E_P_*) using subsets of *D_V_* as they become known. Contrary to the customary model of divide and conquer, where subproblems are completely solved before combining into a global solution, this solves portions of subproblems on each iteration.

**Fig. 4.**
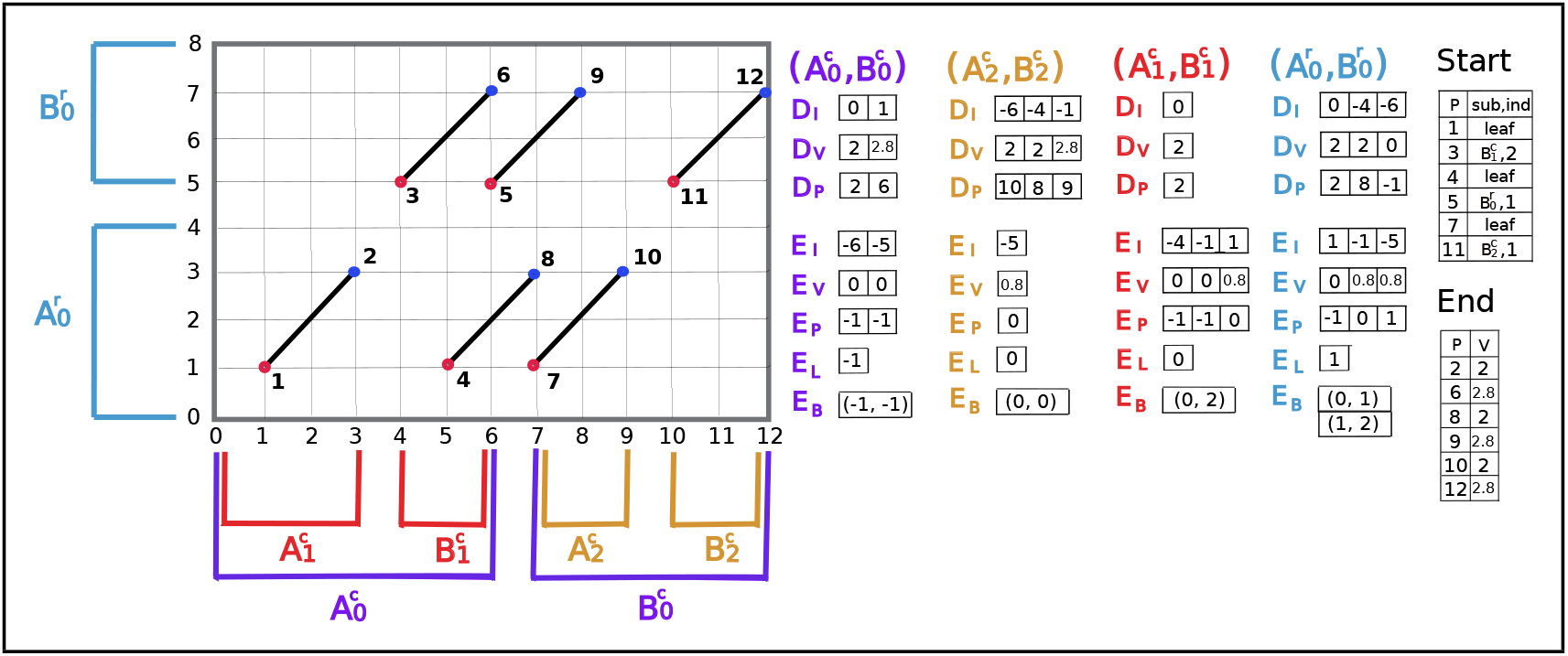
The visualization of subproblems divisions and the data structures needed for each subproblem: *D_I_*, *D_V_*, *D_P_*, *E_I_*, *E_V_*, *E_P_*, *E_L_* and *E_B_* after the processing of all the 12 points. Points are numbered in the order of processing (Cartesian sorted order). 12 points are assigned into three column subproblems 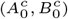, 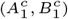, 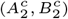 and one row subproblem 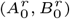, where starting points are assigned to *A*-part and endpoints are assigned to *B*-part. Leaf subproblems are not shown for simplicity. *Start* and *End* are used for the traceback of the optimal chain. *Start* stores *sub* - the index of the subproblem which yields the optimal chaining score up to a starting point and *ind* - *φ*(*E_I_*, *f_i_*), where *f_i_* is the diagonal of the starting point. *End* stores the optimal value for each endpoint. For this toy example, the match bonus of every fragment is 2 and the gap cost function of appending fragment *α_j_* to fragment *α_i_* is 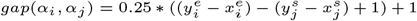, where 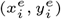 is the endpoint of *α_i_* and 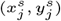 is the starting point of *α_j_*. *End* indicates that there are three optimal chains achieving the optimal value: 2.8. By tracing back, chain-1: point-1, point-2, point-3, point-6, chain-2: point-1, point-2, point-5, point-9 and chain-3: point-7, point-10, point-11, point-12.

Points are processed in order of *x* and then *y* coordinate, determining the value of the optimal chain up to the current processed point. The value of a starting point is the value of the optimal chain that chains the starting point to an endpoint below it. When no endpoints are below a starting point, the value of this starting point is trivially set to 0. The value of an endpoint is simply the value of the optimal chain preceding the corresponding starting point plus the value of the fragment.

When processing an endpoint *p_i_*, the starting point *p_s_* of the corresponding fragment *α_i_* has been solved because points are processed in Cartesian order. The value of the chain at the endpoint is *Score*(*p_s_*) +*l_i_*, where *l_i_* is the match bonus of fragment *α_i_*. In order for *p_i_* to be used when solving for starting points that are above it, the value of *D_V_* must be updated in subproblems for which *p_i_* is a point in the *A*-part. The *SA* list is used to determine which subproblems contain *p_i_* in a *A*-part. Suppose the forward diagonal of *p_i_* is *f_i_*. In each subproblem in SA, *Score*(*p_i_*) is passed to *D_V_* [*j*] and *D_V_* [*j*] is set to max{*Score*(*p_i_*), *D_V_* [*j*]}, where *j* = *φ*(*D_I_, f_i_*), and *D_P_* [*j*] is set to *i* if *D_V_* [*j*] gets updated.

When processing a starting point *p_i_*, the optimal value must be calculated in each of the *B*-parts of subproblems that include *p_i_*, and the global optimal value will be selected among them. For any subproblem, this can be achieved by solving for *E_V_* [*j*] using Equation (4) for column-based or Equation (5) for row-based subproblems, where *j* = *φ*(*E_I_, f_i_*) and *f_i_* is the diagonal of *p_i_*. For a column-based subproblem, this requires that the values of *D_V_* [*k*] have been solved where *D_I_* [*k*] ≤ *f_i_*, and for row-based subproblems, the values of *D_V_* [*k*] must have been solved where *D_I_* [*k*] > *f_i_*. These correspond diagonals overlapping points that fall in the region below and to the left of the *p_i_* and are exactly what have been processed when solving for points in Cartesian sorted order. Thus all required values in *D_V_* arrays (though not the entire *D_V_* array) are available when solving for *p_i_*.

The value of *E_V_* [*j*] is optimal once *Update*(*D_V_* [*k*], *E_B_*) has been called on all diagonals that contain potential predecessors to points on diagonal *E_I_* [*j*]. The function *Update*(*D_V_* [*k*], *E_B_*) must be called only once per element in *D_V_* and in increasing order. However because points are processed in Cartesian sorted order, values of *E_V_* are solved in arbitrary order, and calling *Update*(*D_V_* [*k*], *E_B_*) can reference elements in *D_V_* multiple times. To account for this, we maintain an index *E_L_* for each subproblem that keeps track of the last element of *D_V_* which has been processed by *Update*(*D_V_* [*k*], *E_B_*). Before solving any points, *E_L_* is initialized to −1 in every subproblem, and when processing points will be assigned to reference the most recently updated diagonal from *D_V_*. When processing starting point *p_i_* in a column-based subproblem, *E_V_* [*j*] must be solved, where *j* = *φ*(*E_I_, f_i_*).If *D_I_*[*E_L_*] > *f_i_*, *Update* has been called on all values of *D_V_* that *E_V_* [*j*] relies on, and the state of *E_B_* contains the optimal value for *E_V_* [*j*] such that it may be calculated immediately from Ω(*E_B_, j*). Otherwise, *Update*(*D_V_*[*k*], *E_B_*) iscalledfor *E_L_* < *k* < *C*, where *D_I_* [*C*] is the first diagonal from the left that is larger than *f_i_* in array *D_I_*, and *E_V_* [*j*] may be calculated from Ω(*E_B_, j*). Similarly, for row-based subproblems if *f_i_* ≥ *D_I_*[*E_L_*], the value of *E_V_* [*j*] may be assigned immediately from Ω(*E_B_, j*) where *j* = *φ*(*E_I_, f_i_*). Otherwise, *Update*(*D_V_* [*k*], *E_B_*) is called for *E_L_* < *k* < *C*, where *D_I_*[*C*] is the first diagonal from the left that is smaller than or equal to *f_i_* in array *D_I_*, and *E_V_*[*j*] can be retrieved from Ω(*E_B_, j*). By comparing values *E_V_* [*j*] from all the subproblems in the *SB* list of startpoint *p_i_*, the maximum value for pi will be obtained and stored. The pseudocode and detailed example of solving points and conquering subproblems are given in the supplementary material (Supplementary Algorithm S1 and Figure S2).

#### 3.6.3 Extension to Inversion Cases

This extension is inspired by the work (Brudno *et al.*, 2003). As explained in the previous section, when chaining fragments in forward direction, two points - a lower left start point 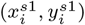 and a upper right endpoint 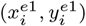 would be associated to each fragment *α_i_*. 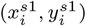 can be chained to an endpoint below and to the left of it and 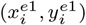 can be the predecessor of a starting point above and to the right of it. To allow inversions to happen, fragments must be allowed to be chained in the reverse direction. To account for this, we associate two more points to each fragment *α_i_*, that are a upper left start point 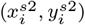 and a lower right endpoint 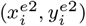. The start point 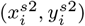 can be only be chained to endpoint 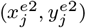 of some other fragment *α_j_* that satisfies 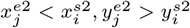. The endpoint 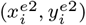 can precede a starting point 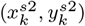 of some other fragment *α_k_* that satisfies 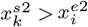, 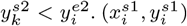, 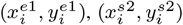 and 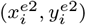 can be represented as *s*1(*i*), *e*1(*i*), *s*2(*i*) and *e*2(*i*) respectively. Chaining fragment *α_i_* to fragment *α_j_* in the reverse direction equals to chaining the start point *s*2(*α_i_*) to the endpoint *e*2(*α_j_*). The cost of appending *s*2(*α_i_*) to *s*2(*α_j_*) in the reverse direction is *gap^rev^*(*α_j_, α_i_*), which is a convex function of 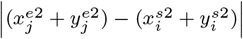, the difference between the reverse diagonals of fragments *α_i_* and *α_j_*. In a short word, *s*2(*α*) and *e*2(*α*) of each fragment *α* would be responsible for the possible chaining of *α* in the reverse direction, while *s*1(*α*) and *e*1(*α*) would be responsible for the forward direction.

We used the same column and row subproblem dividing scheme to sort all *s*1 and *e*1 points and assign them into column-1 and row-1 subproblems. Then all *s*2 and *e*2 points are sorted and assigned to column-2 and row-2 subproblems in the same way. Arrays *D_I_*, *D_V_*, *D_P_*, *E_I_*, *E_V_*, *E_P_*, the block structure *E_B_* and variable *E_L_* are allocated and initialized for each subproblem. Lists *SA*_1_ and *SA*_2_ reference column-1/row-1 and column-2/row-2 subproblems that each endpoint *e*1 and *e*2 is in respectively. Similarly, lists *SB*_1_ and *SB*_2_ references column-1/row-1 and column-2/row-2 subproblems that starting points *s*1 and *s*2 in respectively. The only difference between column-2/row-2 and column-1/row-1 subproblems is that column-2/row-2 stores reverse diagonal instead of forward diagonal in *D_I_* and *E_I_* arrays.

The steps to solve subproblems to allow chaining in both forward and reverse directions are highly similar to section 3.6.2. Points are processed in order of *x* and then *y* coordinate. When a starting point *s*1(*α_i_*) is being processed, *E_V_*[*j*], where *j* = *φ*(*E_I_, f_i_*) and *f_i_* is the forward diagonal, will be computed from column-1/row-1 subproblems in *SB*_1_. The maximum *E_V_* [*j*] of those subproblems is the value of optimal chain up to *s*1(*α_i_*), *Score*(*s*1(*α_i_*)), where *s*1(*α_i_*) is forwardly chained to an endpoint *e*1(*α_j_*) below and to the left of it. Similarly, when a starting point *s*2(*α_i_*) is being processed, *E_V_*[*j*], where *j* = *φ*(*E_I_, r_i_*) and *r_i_* is the reverse diagonal, will be computed from column-2/row-2 subproblems in *SB*_2_. The maximum *E_V_*[*j*] of those subproblems is the value of optimal chain up to *s*2(*α_i_*), *Score*(*s*2(*α_i_*)), where *s*2(*α_i_*) is reversely chained to an endpoint *e*2(*α_k_*) above and to the left of it.

After solving *s*1(*α_i_*) and *s*2(*α_i_*) for fragment *α_i_*, the value of the optimal chain up to fragment *α_i_* can be calculated as *Score*(*α_i_*) = max{*Score*(*s*1(*α_i_*)), *Score*(*s*2(*α_i_*))} + *l*(*α_i_*), where *l*(*α_i_*) is the match bonus of fragment *α_i_*. This optimal chain is chosen from all the possible chains that fragment *α_i_* is forwardly or reversely chained to the predecessor in the left. When solving *e*1(*α_i_*), *Score*(*α_i_*), will be passed to array *D_V_*[*j*], where *j* = *φ*(*E_I_, f_i_*) and *f_i_* is the forward diagonal, to column-1/row-1 subproblems in *SA*_1_. And *D_P_* [*j*] will be updated to the index of point *e*1(*α_i_*), if *Score*(*α_i_*) > *D_V_*[*j*]. Similarly, when *e*2(*α_i_*) is being processed, *Score*(*α_i_*) will be passed to array *D_V_* of column-2/row-2 subproblems in *SA*2 and *D_P_* will be updated.

Therefore, the addition of two points *s*2(*α*) and *e*2(*α*) for each fragment *α* make it possible to allow *α* to be chained in the reverse direction. Meanwhile, the overall time complexity and storage remain bounded by *O*(*n*(log(*n*)^2^)), where *n* is the total number of fragments.

#### 3.6.4 Time Complexity

Assume there are n fragments in total, list *SB/SA* contains *O*(log(*n*)) subproblems that a point is in. In section 3.6.2, we mention that when a starting point pi with forward diagonal *f_i_* is being processed, *Update*(*D_V_* [*k*], *E_B_*) is called for *E_L_* < *k* < *C*, where *D_I_* [*C*] is the first diagonal from the left that is larger than/smaller than or equal to *f_i_* in column/row subproblems. And *E_V_* [*j*] can be retrieved in *O*(log(*n*)) time from the block structure *E_B_* by calling Ω(*E_B_, j*), where *j* = φ(*E_I_*, *f_i_*). Procedure Update may be called several times for a starting point in each subproblem. In order to make it easy to quantify the total time complexity of Update procedures, we consider that each Update procedure is called right after *D_V_* [*j*] is updated by some endpoint *p_i_* with diagonal *f_i_*, where *j* = φ(*E_I_, f_i_*). When an endpoint is being solved, there are *O*(log(*n*)) subproblems associated with it and in each subproblem *Update* takes O(log(*n*)) to conduct. Therefore, it takes O((log(*n*))^2^) time to solve the subproblems that are associated with an endpoint. When a starting point is being processed, there are *O*(log(*n*)) subproblems it is in and in each subproblem *E_V_*[*j*] can be computed from the block structure *E_B_* in O(log(*n*)) time. Therefore, it takes O((log(*n*))^2^) time to tackle subproblems that are associated with a starting point. Since there are n fragments in total, the time complexity of processing all the points and subproblems is bounded by O(n log(n)^2^).

## 4 Discussion

The initial description of CG-SDP was given in 1992 in two publications, one covering an affine-cost gap function, and another with a convex-cost gap function.(Eppstein *et al.*, 1992a,b). While there have been many implementations of affine cost SDP, no sequence alignment methods have been implemented using SDP with a convex-cost gap function. In the original description of the algorithm, the processing of a starting point is blocked until all subproblems the point relies on are solved, and then the process is unblocked and processing resumes. We find that this blocking and unblocking strategy is not necessary, and the addition of data structures to keep track of the state of computation of subproblems enables solving the problem with a standard model of computation. We used two additional strategies to effectively employ SDP in lra: an iterative refinement process where a large number of anchors from the initial minimizer search are grouped into a small super-fragments that are chained using SDP, and once a rough alignment has been found a new set of matches with smaller anchors is calculated using the local miminizer indexes. As a result, alignment is of similar speed to state of the art algorithms, without the need for single-instruction multiple-data (SIMD) processing; runtime was slower when using an SIMD alignment library (Šošic and Šikic, 2017) possibly due to the overhead of invoking the library functions.

The lra alignments have competitive runtime and memory usage compared to minimap2. Using two different SV discovery algorithms, pbsv and Sniffles, we show it is possible to use lra alignments to discover SV using PacBio HiFi, CLR, and Oxford Nanopore reads, as well as directly from aligned *de novo* assembly contigs. The performance for SV detection using PacBio reads and the pbsv algorithm is similar between lra and minimap2, with lra demonstrating a small gain in recall over larger SV events. Importantly, the availability of multiple alignment algorithms can help improve the results of studies that require high sensitivity and specificity, such as Mendelian analysis. The greater improvement in SV discovery metrics on ONT data aligned by lra additionally highlights the utility of using multiple algorithms to analyze sequencing data.

Finally, as *de novo* assembly of genomes becomes more routine, it is important to have accurate methods of SV detection from contig alignments. We used lra to align a haplotype-resolved genome assembled from PacBio HiFi reads and detect SV. When compared to the GIAB SV benchmark data, the lra alignments show slightly lower precision and recall than read-based SV detection. However, when the assembly-based SV are compared to SV detected in an orthogonal read dataset, nearly all (>98.7%) variants discovered from assemblies are supported by read alignments. This indicates few differences between the HiFi based assembly and the reference are due to assembly error, and the annotated precision of the callset is likely an under estimate. A variant callset is effectively a list of operations that may be applied to the reference genome to reconstruct a sample genome. Because the HiFi assembly has few assembly errors on the same size scale as an SV, this supports the development of an alternative model for validating SV callsets in which the reconstructed genome is compared to the haplotype-resolved assembly, rather than by comparing callsets. This may be used to validate calls outside the high-confidence regions defined in In regions that a haplotype-resolved assembly has been confidently generated

## 5 Acknowledgments

J.R. is supported by the Vitterbi Fellowship. M.J.P.C. is supported by NHGRI U24HG007497.

## 6 Data Availability

The HG002 assembly data is available in NCBI database under GenBank assembly accession number GCA_004796285.1 and GCA_004796485.2. The HG002 ONT reads are available in NCBI database under BioProject accession number PRJNA678534. The HG002 HiFi and CLR reads can be downloaded using links supplied in the supplement. Variant calls, and a set of curated inversions are available at https://figshare.com/articles/dataset/lra-supplemental-HG002-SV_vcf_tar_gz/13238717.

