## Supplementary Materials for "lra: the Long Read Aligner for Sequences and Contigs"

### Supplementary Material

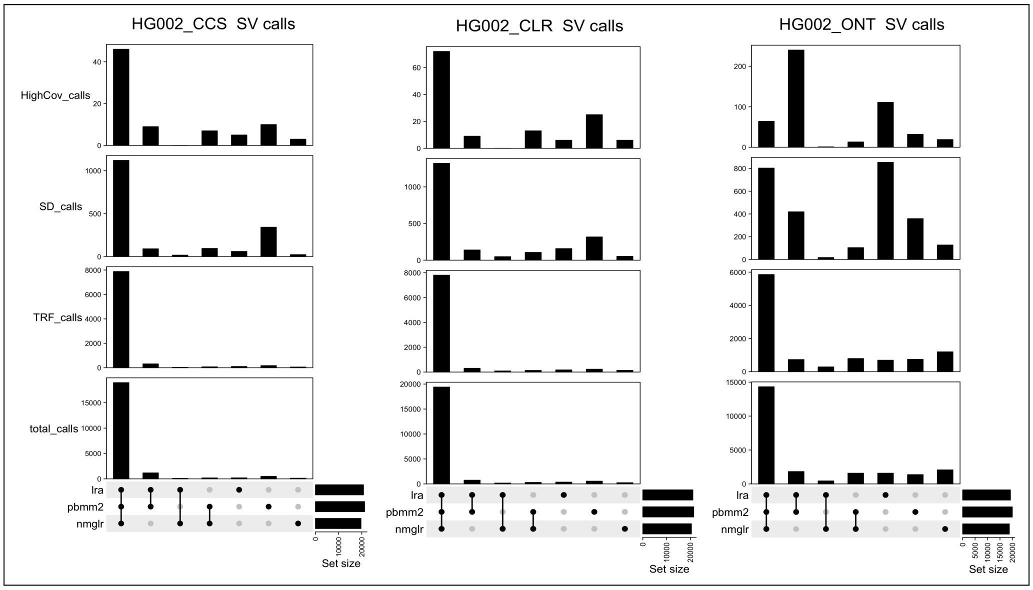


**Fig. S1.** Intersections between the SV callsets from every two aligners and aligner-unique calls for HG002 CCS, CLR and ONT datasets. It further displays locations annotations of shared/aligner-unique calls – residing in segmental duplications/tandem repeats and annotates calls with coverage higher than 2*genome wide coverage.

#### 1. DATA PATHS

s3-us-west-2.amazonaws.com/human-pangenomics/HG002/hpp$_{H}G002_{N}A24385_{s}on_{v}1/PacBio_{H}iFi/19kb/m64011_{1}90714_{1}2074$
$us-west-2.amazonaws.com/human-pangenomics/HG002/hpp_{H}G002_{N}A24385_{s}on_{v}1/PacBio_{H}iFi/19kb/m64011_{1}90728_{1}112$
$us-west-2.amazonaws.com/human-pangenomics/HG002/hpp_{H}G002_{N}A24385_{s}on_{v}1/PacBio_{H}iFi/20kb/m64011_{1}90830_{2}201$
$us-west-2.amazonaws.com/human-pangenomics/HG002/hpp_{H}G002_{N}A24385_{s}on_{v}1/PacBio_{H}iFi/20kb/m64011_{1}90901_{0}953$
$us-west-2.amazonaws.com/human-pangenomics/HG002/hpp_{H}G002_{N}A24385_{s}on_{v}1/PacBio_{H}iFi/25kb/m64011_{1}90712_{2}257$
$us-west-2.amazonaws.com/human-pangenomics/HG002/hpp_{H}G002_{N}A24385_{s}on_{v}1/PacBio_{H}iFi/25kb/m64011_{1}90726_{2}203$

The HG002 CLR reads can be downloaded using the following links:

s3-us-west-2.amazonaws.com/human-pangenomics/HG002/hpp$_{H}G002_{N}A24385_{s}on_{v}1/PacBio_{C}LR/WUSTL_{S}V-HG002-CLR/1_{A}01/m64043_{1}91010_{1}74437.subreads.bams3-us-west-2.amazonaws.com/human-pangenomics/HG002/hpp_{H}G002_{N}A24385_{s}on_{v}1/PacBio_{C}LR/WUSTL_{S}V-HG002-CLR/3_{C}01/m64043_{1}91012_{1}02127.subread$

**Algorithm S1.** Defining subproblems

---

**Require:** $X$ - the sorted set of starting and ending points from the set of anchors $\Phi$; $[s, e]$ - the column/row range; $d \in \{col, row\}$;

**Ensure:** SUB$(d)$ - the desired set of column/row subproblems obtained by assigning points from column/row $s$ to column/row $e$, which is initialized as $\emptyset$;

**procedure** SUB$(d, s, e, X)$
    **if** $e == s$ **then**
        Construct subproblem $(d, \emptyset, \emptyset, s, e, \text{DATA})$ with DATA$= (\emptyset, \emptyset, \emptyset, E_I, E_P, E_V)$ as following:
        Save the forward diagonals of all the starting points from column/row $e$ in array $E_I$;
        Sort the forward diagonals in array $E_I$ in the increasing/decreasing order;
        Initialize every entry in $E_V$ to 0;
        Initialize every entry in $E_P$ to -1;
        SUB$(d) \leftarrow$SUB$(d) \cup \{(d, \emptyset, \emptyset, s, e, \text{DATA})\}$;
    **else**
        **while** $e > s$ **do**
            Construct subproblem $(d, s, \lfloor (s+e)/2 \rfloor, \lfloor (s+e)/2 \rfloor + 1, e, \text{DATA})$ with DATA$= (D_I, D_P, D_V, E_I, E_P, E_V)$ as following:
            Store the forward diagonals of all the ending points from column/row $s$ to column/row $\lfloor (s+e)/2 \rfloor$ in array $D_I$;
            Sort the forward diagonals in array $D_I$ in the increasing/decreasing order;
            Initialize every entry in $D_V$ to 0;
            Initialize every entry in $D_P$ to -1;
            Save the forward diagonals of all the starting points from column/row $\lfloor (s+e)/2 \rfloor + 1$ to column/row $e$ in array $E_I$;
            Sort the forward diagonals in array $E_I$ in the increasing order;
            Initialize every entry in $E_V$ to 0;
            Initialize every entry in $E_P$ to -1;
            SUB$(d) \leftarrow$SUB$(d) \cup \{(d, s, \lfloor (s+e)/2 \rfloor, \lfloor (s+e)/2 \rfloor + 1, e, \text{DATA})\}$;
            SUB$(d, s, \lfloor (s+e)/2 \rfloor, X)$;
            SUB$(d, \lfloor (s+e)/2 \rfloor + 1, e, X)$;
    return SUB$(d)$;

---

**Algorithm S2.** Sparse Dynamic Programming with convex gap cost

Get the points set $X$ from the set of anchors $\Phi$;
Sort $X$ in Cartesian order; Assume all points are arranged from col 0 to col $t$;
SUB$\leftarrow \emptyset$; ▷ SUB is the set storing all subproblems
SUB$\leftarrow$SUB$\cup$SUB$(col, 0, t, X)$;
Sort $X$ in anti-Cartesian order; Assume all points are arranged from row 0 to row $q$;
SUB$\leftarrow$SUB$\cup$SUB$(row, 0, q, X)$;
**for** each $p_i \in CartesianSort(X)$ **do**
  **if** $p_i$ is an endpoint **then**
    $Score(p_i) \leftarrow Score(p_s) + l_i$, where $p_s$ is the corresponding startpoint and $l_i$ is match bonus of the corresponding anchor;
    $End(p_i) \leftarrow Score(p_i)$; ▷ $End$ stores the optimal chaining score for each endpoint
    **for** each SUB[j]$\in SA$ **do**
      **if** $Score(p_i) > D_V[j]$ **then**
        $D_P[j] \leftarrow i$;
        $D_V[j] \leftarrow Score(p_i)$, where $j = \varphi(D_I, f_i)$ and $f_i$ is the forward diagonal of $p_i$;
  **else if** $p_i$ is a starting point **then**
    $maxvalue \leftarrow 0$;
    **for** each SUB$[h] \in SB$ **do**
      $j \leftarrow \varphi(E_I, f_i)$, where $f_i$ is the forward diagonal of $p_i$;
      **if** SUB$[h]$ is a col-based subproblem **then**
        **if** $D_I[E_L] > f_i$ **then**
          $E_V[j] \leftarrow \Omega(E_B, j)$;
        **else**
          **for** each $k \in (E_L, C)$, where $C = \min_s D_I[s] > f_i$ **do**
            $Update(D_V[k], E_B)$;
          $E_L \leftarrow C - 1$;
          $E_V[j] \leftarrow \Omega(E_B, j)$;
      **else if** SUB$[h]$ is a row-based subproblem **then**
        **if** $D_I[E_L] \leq f_i$ **then**
          $E_V[j] \leftarrow \Omega(E_B, j)$;
        **else**
          **for** each $k \in (E_L, C)$, where $C = \min_s D_I[s] \leq f_i$ **do**
            $Update(D_V[k], E_B)$;
          $E_L \leftarrow C - 1$;
          $E_V[j] \leftarrow \Omega(E_B, j)$;
      **if** $E_V[j] > maxvalue$ **then**
        $Start[p_i] \leftarrow ($SUB$[h], j)$

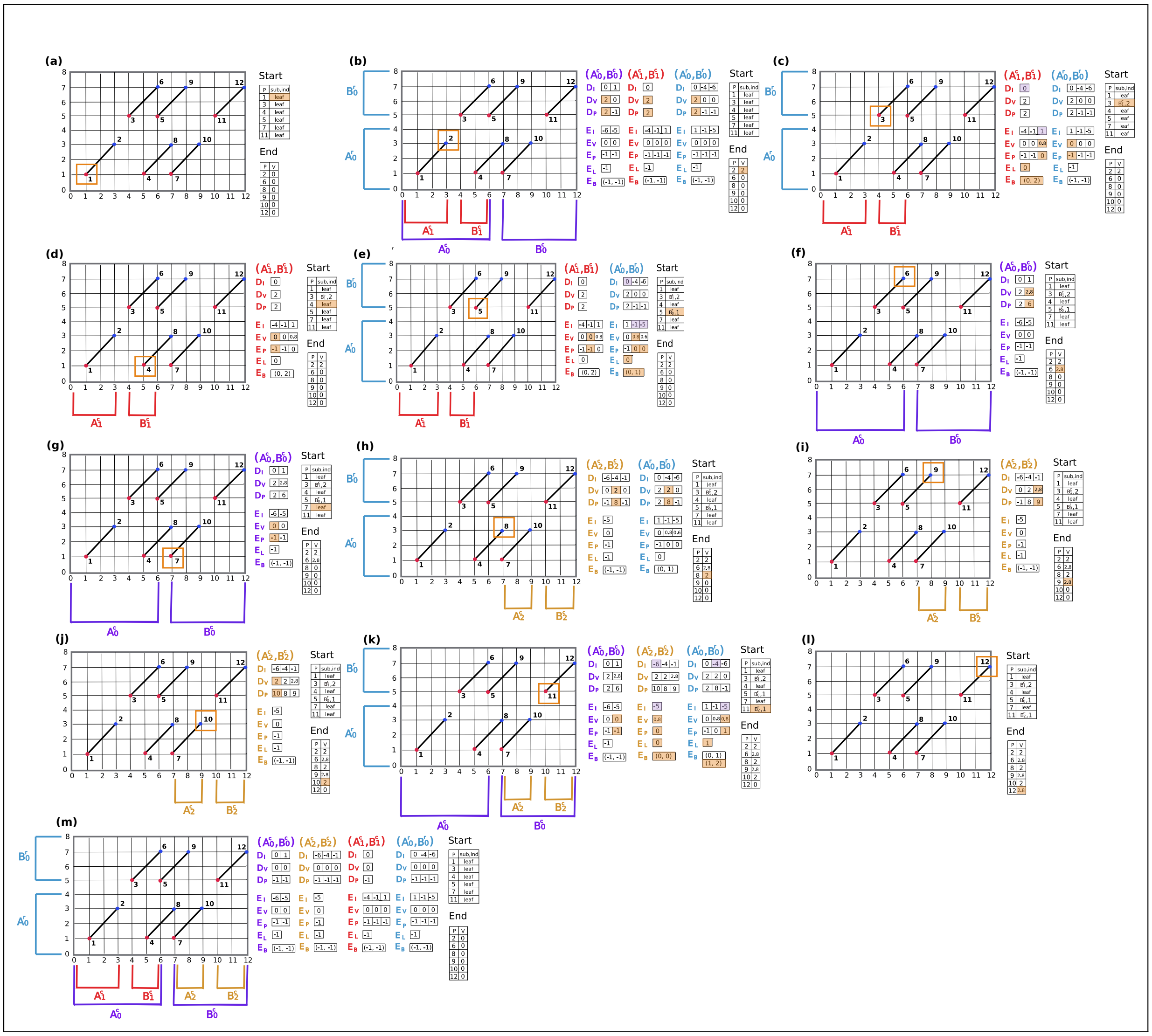

**Fig. S2** ***(previous page).*** A detailed example of visualizing subproblems division, the data structures for each subproblem: $D_I$, $D_V$, $D_P$, $E_I$, $E_V$, $E_P$, $E_L$, $E_B$ and the process of subproblems solving. The horizontal axis represents the query, while the vertical axis represents the target. Points are numbered in Cartesian sorted order, which is the processing order. 12 points are assigned into three column subproblems $(A_0^c, B_0^c)$, $(A_1^c, B_1^c)$, $(A_2^c, B_2^c)$ and one row subproblem $(A_0^r, B_0^r)$, where starting points are assigned to $A$-part and endpoints are assigned to $B$-part. Leaf subproblems are not shown for simplicity. Start and End are used for the trace-back of the optimal chain. $Start$ stores $sub$ – the index of the subproblem which yields the optimal chaining score up to a starting point and $ind$ – the index of $f_i$ in array $E_I$, that is $\varphi(E_I, f_i)$, where $f_i$ is the diagonal of the starting point. $End$ stores the optimal value for each endpoint. For this toy example, gap cost of appending fragment $\alpha_j$ to fragment $\alpha_i$ is $G(\alpha_i, \alpha_j) = 0.25 * \log((y_i^e - x_i^e) - (y_j^s - x_j^s) + 1) + 1$, where $(x_i^e, y_i^e)$ is the endpoint of $\alpha_i$ and $(x_j^s, y_j^s)$ is the startpoint of $\alpha_j$. **m**, shows the regions that each subproblem covers and the initialized data structures for each subproblem. There are of three column subproblems and one row subproblems (leaf subproblems are not shown for simplicity): $(A_0^c, B_0^c)$, $(A_1^c, B_1^c)$, $(A_2^c, B_2^c)$ and $(A_0^r, B_0^r)$. **a-l** shows how the data structures of subproblems that are associated with the point being processed in each step are updated. Note that entries that are updated are highlighted by orange. **a**, shows for $startpoint - 1$, it is a leaf subproblem that yields the value of the optimal chain up to $startpoint - 1$. **b**, shows when processing $endpoint - 2$, the optimal value up to it is $Score(startpoint - 1) + 2$, where 2 is the match bonus of the fragment. Array $End$ is then updated. Since $endpoint - 2$ is in $A_0^c$, $A_1^c$, $A_0^r$, the corresponding $D_V$ entries are updated to 2 and corresponding $D_P$ entries update to the index of $endpoint - 2$. **c**, shows when processing $startpoint - 3$, it is in the $B$-parts of subproblems $(A_1^c, B_1^c)$ and $(A_0^r, B_0^r)$. $startpoint - 3$ is located in $E_I[2]$ of $(A_1^c, B_1^c)$, so $Update(D_V[0], E_B)$ would be called to get the value of $E_V[2]$. The purple color highlighting shows what forward diagonals in $E_V$ would be updated by $D_V[0]$. $E_P[2]$ would be updated to point to $D_I[0]$. $startpoint - 3$ is located in $E_I[2]$ of $(A_0^r, B_0^r)$ and no forward diagonals in $D_I$ used to update $E_V[2]$. Therefore, in $Start$, $sub$ and $ind$ for $startpoint - 3$ are updated to $B_1^c, 2$. **d**, shows when processing $startpoint - 4$, it is in $B_1^c$ and locates in $E_I[0]$ of $(A_1^c, B_1^c)$. Since there is no forward diagonal in $D_I$ can be used to update $E_V[0]$, it is a leaf subproblem that yields the optimal chaining value up to $startpoint - 4$ in $Start$. **e**, shows when processing $startpoint - 5$, it is in $B_1^c$ and $B_0^r$. In $(A_1^c, B_1^c)$, there is no forward diagonal can be used to update $E_V[1]$. In $(A_0^r, B_0^r)$, $Update(D_V[0], E_B)$ is called to update the block structure $E_B$, so $E_V[1]$ and $E_V[2]$ would be computed from $E_B$. In $Start$, $sub$ and $ind$ for $startpoint - 5$ are updated to $B_0^r, 1$. **f**, **g**, **h**, **i**, **j**, **k**, **l**: show the subproblems solving and data structures updating for the rest of points. After processing all 12 points, three optimal chains can be obtained by tracing back, which are $chain - 1 = [startpoint - 1, endpoint - 2, startpoint - 3, endpoint - 6]$, $chain - 2 = [startpoint - 1, endpoint - 2, startpoint - 5, endpoint - 9]$ and $chain - 3 = [startpoint - 7, endpoint - 10, startpoint - 11, endpoint - 12]$.

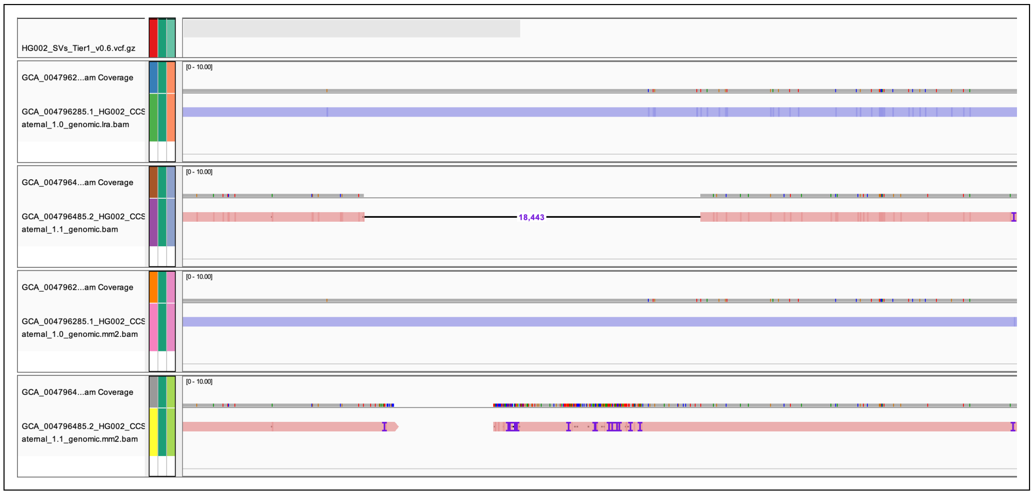


**Fig. S3.** The top track shows the annotated HG002 SV set with high confidence. The second and third tracks are lra alignments of HG002 paternal and maternal assemblies and the rest two are minimap2 alignments. The third track shows a clear insertion of 18443 base pairs in lra alignment of maternal assembly, while the fifth track shows lots of indels clustered around the insertion in the minimap2 alignment.

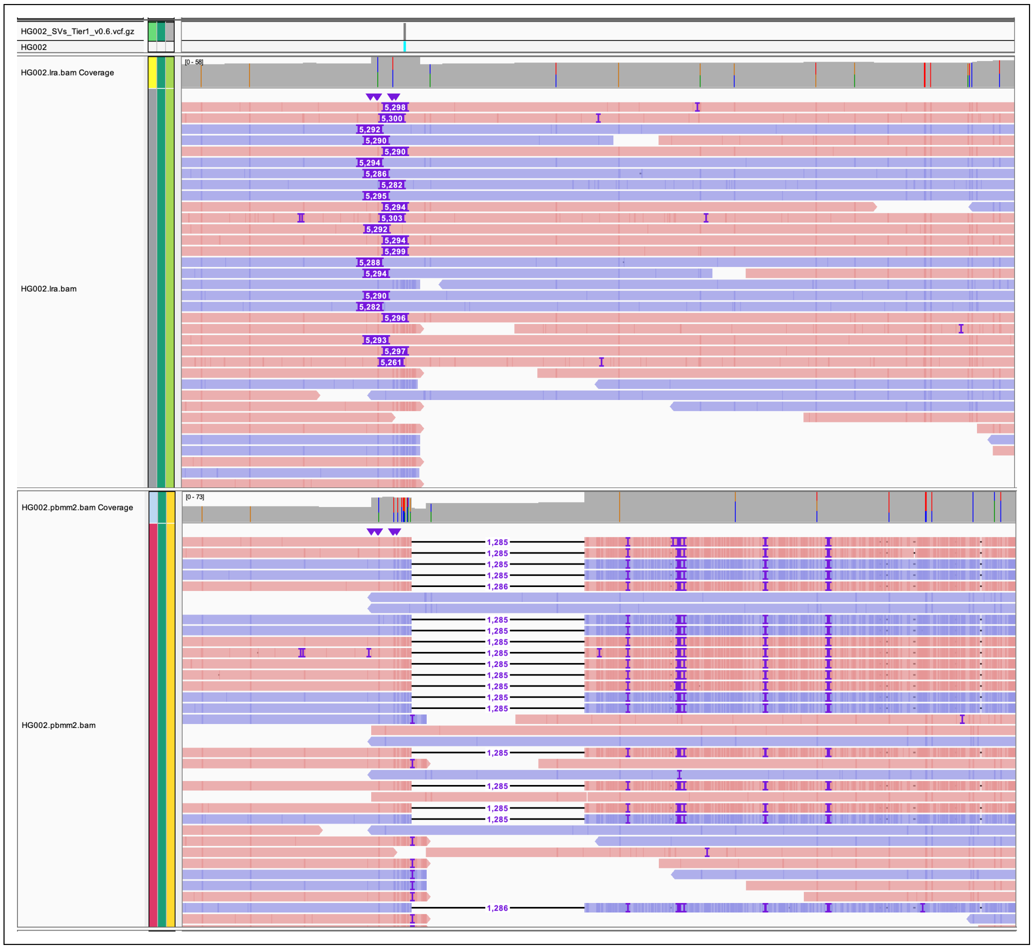


**Fig. S4.** The top track shows the annotated HG002 SV set with high confidence. The second track is lra alignment of HG002 CCS reads and the third one is pbmm2 alignment. There is a clear 5.3k insertion in lra alignment, which also appears in the annotated HG002 SV set with high confidence. However, pbmm2 alignment shows a deletion of 1285 base pairs surrounded by lots of indels at the same location.

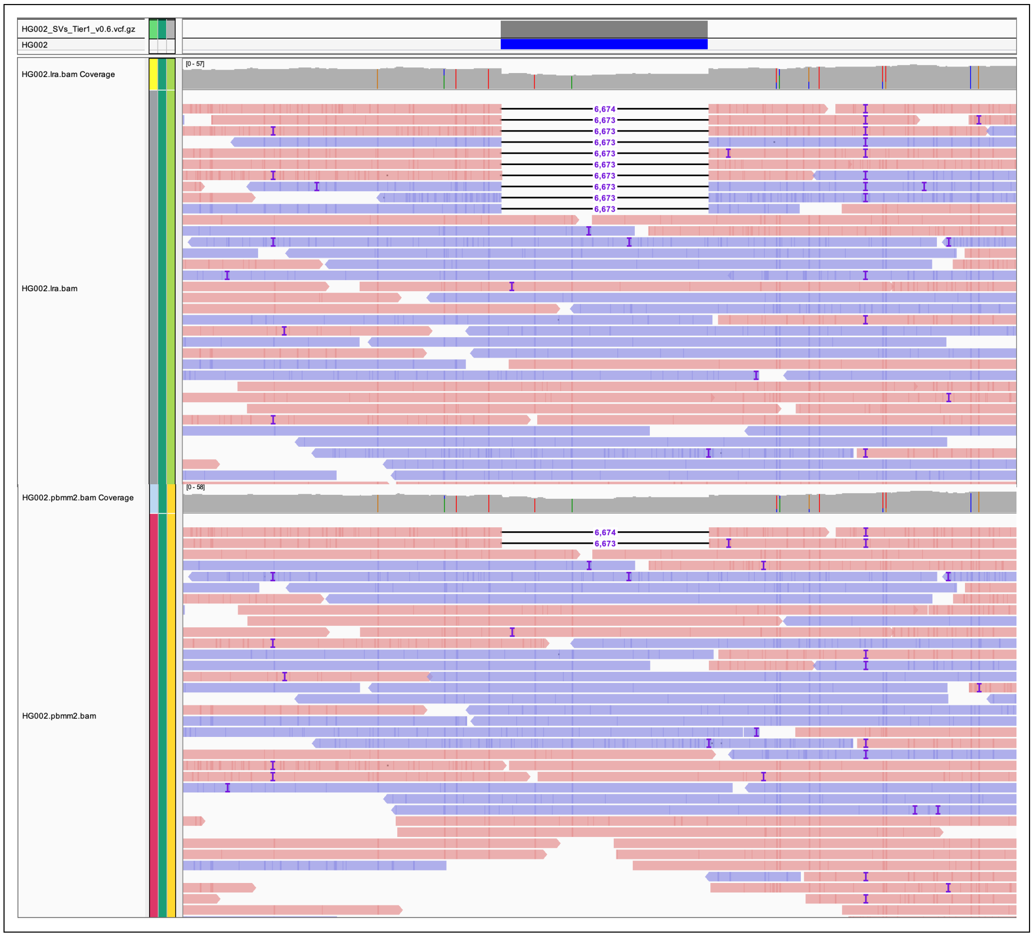


**Fig. S5.** The top track shows the annotated HG002 SV set with high confidence. The second track is lra alignment of HG002 CCS reads and the third track is pbmm2 alignment. The figure shows that lra alignment shows more support for the deletion of 6674 base pairs than pbmm2 alignment.

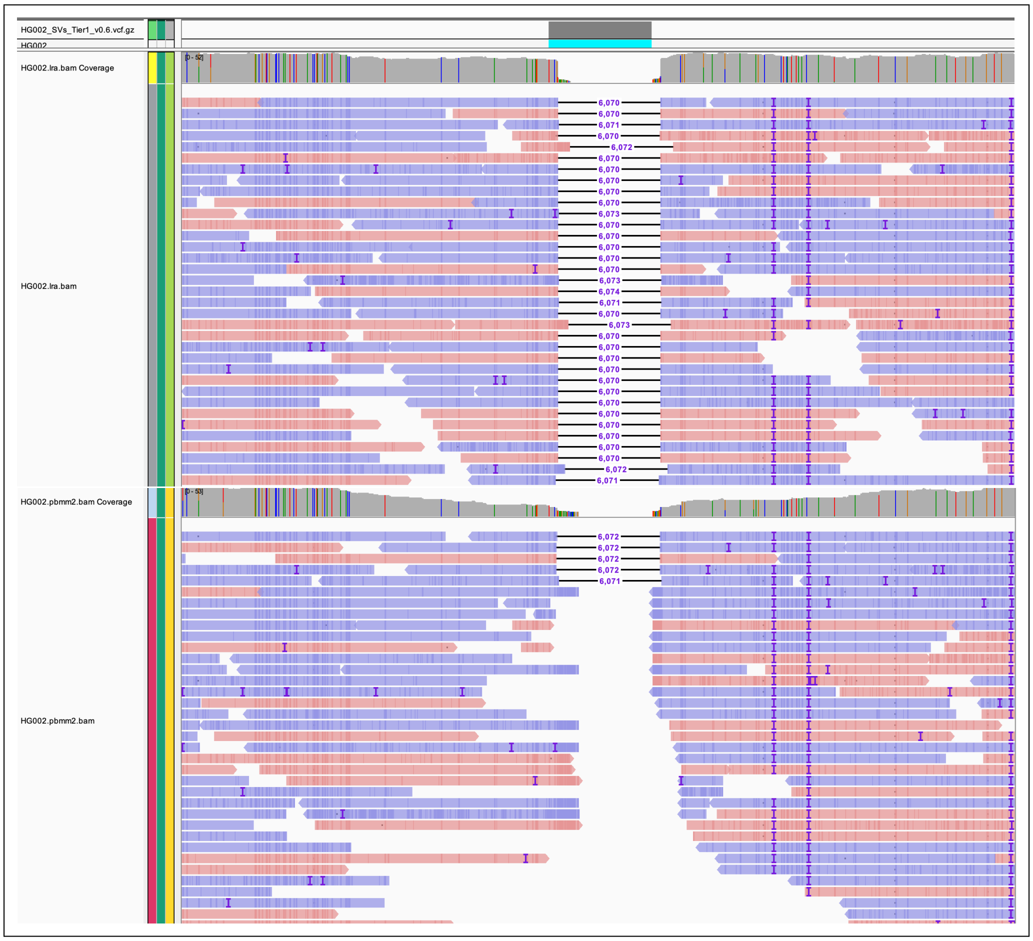


**Fig. S6.** The top track shows the annotated HG002 SV set with high confidence. The second track is lra alignment of HG002 CCS reads and the third track is pbmm2 alignment. The figure shows that lra alignment shows more support for the deletion of 6070 base pairs than pbmm2 alignment.
